# Microarthropod contributions to fitness variation in the common moss *Ceratodon purpureus*

**DOI:** 10.1101/2020.12.02.408872

**Authors:** Erin E. Shortlidge, Adam C. Payton, Sarah B. Carey, Stuart F. McDaniel, Todd N. Rosenstiel, Sarah M. Eppley

**Affiliations:** Portland State University, Department of Biology, P.O. Box 751, Portland, Oregon, 97202-0751, USA; University of Florida, Department of Biology, P.O. Box 118525, Gainesville, Florida, 32611-8525, USA

**Keywords:** bryophyte, *Ceratodon purpureus*, dioecious, fitness, fertilization, mating system, microarthropods

## Abstract

The evolution of mutualism depends critically upon genetic variation in the fitness benefit to both partners. Estimates of these quantities are rare, however, because genetic variation for the interaction may be absent, aspects of the interaction may not be amenable to experimental manipulation, or the benefits to one partner may be unknown. In vitro experiments show that female mosses produce odors which attract sperm-dispersing microarthropods, but the fitness consequences of this interaction for either partner are unknown. Here we established experimental mesocosms to test for a commensal effect of sperm-dispersing microarthropods on moss reproduction. We found that of moss grown with microarthropods showed increased moss reproductive rates by five times, relative to control mesocosms, but remarkably also increased the number of reproducing genotypes, and changed the rank-order of fitness for both male and female genotypes. These results provide an estimate of the fitness benefit for mosses in the presence of microarthropods, and highlight the potential for biotic dispersal agents to alter fitness among moss genotypes in this relationship.

## Background

Mutualism, and facilitative interactions in general, are ubiquitous in nature. Animal-mediated fertilization in plants likely arose as early as the Devonian in non-vascular plants [1], cycads in the Triassic [2] and gymnosperms, and angiosperms in the Cretaceous [3]. Nevertheless, the fitness consequences of such syndromes can be difficult to study because genetic variation for the interaction is absent or aspects of the interaction may be experimentally intractable. In angiosperms, the fitness effects of biotic pollinators are well-examined [4–6]; however, in mosses, for which there is some evidence of animal-mediated fertilization, little is known about how animal-plant interactions influence fitness.

Gene flow shapes patterns of genetic variation within individuals and among populations, and may influence long-term evolutionary trajectory of a lineage [7–9]. Sessile organisms employ a variety of strategies to promote outcrossing and gene flow. Many angiosperms rely on biotic agents to disperse pollen from one plant to another [7, 10–12], which assures seed production, and can promote outcrossing while reducing the number of gametes lost in interspecies mating [e.g., 13, 14–18]. The pollinators in return gain resources themselves [4–6]. Animals may also disperse gametes in other plant groups, such as ferns and bryophytes. Although these plants release water-dispersed motile sperm [19–23], naturally occurring microarthropods [1, 24] such as Oribatid mites and common springtails, (Collembolan species *Folsomia Candida* and *Sinella curviseta*), can enhance sexual reproduction (i.e., sporophyte formation) in laboratory moss cultures [25],[26]. Remarkably, springtails also choose female mosses over male mosses in olfactory choice tests [38]. These observations suggest that mosses and microarthropods participate in scent-mediated fertilization syndrome, much like angiosperms and their pollinators. Yet, we know little about the influence of this commensal interaction on the dispersal of moss gametes or in the resulting the genetic variation of mosses under natural conditions [27, 28].

Here we used custom-constructed outdoor experimental mesocosms to test if the presence of moss-dwelling microarthropods increased reproduction of the moss species *Ceratodon purpureus* [29–33]. We hypothesized that microarthropods facilitate reproduction by dispersing moss sperm both further and more directly to female archegonia.

## Materials and Methods

### Study populations

We collected *C. purpureus* gametophytes from three populations in and near Portland, OR, USA. We air-dried samples from each population and isolated single gametophytes for further study. We identified the sex of each gametophyte based on the presence of male or female gametangia. Each gametophyte was finely ground, and plant fragments were used to cultivate protonema. This process was repeated until many of the same individuals were growing simultaneously in the greenhouse. All plants received the same environmental conditions. The starter cultures were grown in the greenhouse for 24 months before creating the experimental moss mesocosms (see below).

### Mesocosm establishment and cultivation

Sixteen 38-liter pots were filled with a blend of commercial sand and peat moss (2:1); upon examination under a dissecting microscope, the substrate contained no discernable microarthropods. The pots provided adequate buffering from excessive cold and drought (E. Shortlidge, unpublished results). The mosses were applied as solutions of homogenous tissue. For all sixteen mesocosms, the female moss solution was the same. This solution consisted of nine female genotypes, 4 g each, combined to a total 36 g of female moss tissue (F1-F9). The tissue was sifted and homogenized in 25 ml tap water in small batches. The blended moss-water combination was then divided into sixteen 50 mL Falcon tubes of female solution, one per mesocosm.

Two different male solutions were used (Males A and B), each with three genotypes (for a total of six male genotypes in the experiment; M1-M6); 4 g of each male tissue was combined, mixed and homogenized in small batches, resulting in two 50 mL Falcon tubes, each containing 12 g of mixed male tissue, either Males A or B.

We designed the mesocosms with the male mosses growing in the center of the pots, and female mosses surrounding the males. A 10.5 cm diameter plastic disc was placed in the center of each pot, covering the center, while the female solution was applied to the surrounding, uncovered soil surfaces using a syringe. Upon removal of the disc, one of the two male solutions (either Males A or B) was then applied to the center of the pot. Male concentration was higher than that of females per area by about 2x.

Mesocosms were initially kept in the greenhouse and misted twice daily for two months. Mosses were kept at 18°C and a fourteen-hour photoperiod of ~200 μE and a nighttime temperature of 10°C. Each mesocosm was fit with a translucent Open Top Chamber (OTC) ring that transmits full spectrum sunlight. The OTC rings served as a barrier to prevent excess invertebrate immigration or emigration and to assist in providing uniform canopy temperatures across mesocosms.

Two months later (February 2013), the mesocosms had grown into uniform, yet still short, mats of *C. purpureus* gametophytes (<3 cm tall). The mesocosms were moved outside and except for occasional supplemental misting, the plants grew in natural outdoor conditions, including at least one winter freeze and snowfall event.

### Microarthropod additions

To test the effects of microarthropods on moss reproduction, microarthropods were added to half of the mesocosms. Microarthropods were sourced from naturally occurring mats of mosses (largely *C. purpureus*) found and collected near Portland, OR. Collected moss mats were uniformly misted with tap water, weighed into 100 g portions, and added to modified, collapsible Berlese funnels for live invertebrate extraction under 15W incandescent bulbs [27, 34–36]. Over a year, nine 48-hour live microarthropod extractions were conducted. A tent containing nine suspended extraction funnels was placed over the mesocosms, allowing for live microarthropod extractions to occur without moving the mesocosms. One control extraction was conducted during each extraction time, allowing us to quantify the general abundance and composition of invertebrate additions. These resulted in an average of 356 (± 106 SD) invertebrates per extraction. The extractions were largely comprised of Collembolans and mites (Oribatida and Prostigmata), as well as other invertebrates including species of: Annelida, Arachnida, Coleoptera, Diptera, Hymenoptera, and Nematoda. Post-extraction, the dried mosses were returned to the local environment.

### Canopy physiology

In February 2014, canopy physiology measures were made on leafy gametophytic moss cover. Sporophytes had begun to develop in seven of 16 mesocosms. We determined mesocosm moss canopy chlorophyll content by chlorophyll fluorescence [37] (reported as CFR, chlorophyll fluorescence ratio), using a hand-held meter (Opti-Sciences, CCM-300 Chlorophyll Content Meter, Hudson NH, USA), using standard manufacturer recommended protocols, five values per mesocosm were averaged to obtain one data point per mesocosm. In addition, to non-invasively assess chlorophyll fluorescence parameters, we measured maximum quantum yield of PSII (Fv/Fm). Fluorescence was measured on dark-adapted *C. purpureus* at five locations in each mesocosm before sunrise [38, 39]. In May, we repeated the chlorophyll content measures.

### Sporophyte collection and location

In May 2014, after 15 months, twelve of sixteen mesocosms had developed mature sporophytes, and we began collecting sporophytes. Each mature sporophyte was surveyed for distance from center and angle vector, and carefully pulled from the mesocosm with forceps - along with its maternal gametophyte when feasible. Each sporophyte’s height was measured with digital calipers, recorded and placed into a labeled conical tube with the adjacent maternal tissue for subsequent genetic analysis.

### Parentage analysis

To gage the parentage of a subset of sporophytes, we used a novel, comprehensive genotyping approach. Spores from each sampled sporophyte were grown in axenic culture for genetic sequencing. The haploid father of each sporophyte was derived through identification of maternal gametophyte and subtraction of that from the attached diploid sporophyte (as determined from spores). Upon subtracting the maternal haploid genotype from the diploid sporophyte genotype, paternity can be derived by comparison of deduced paternal genotype to already genotyped males. Further, the spores from the sporophytes were isolated, counted and germinated providing a direct measure of fitness above and beyond sporophyte production [40].

### Sample culture

In total, 325 operculate sporophytes were surface sterilized for 25 s in a 20% solution of commercial bleach (8.25% sodium hypochlorite) and triple rinsed in sterile distilled water before the spores were released into 1 mL of sterile water by mechanically disrupting the capsule. 10 μL of spore suspension per sporophyte was germinated on standard BCDA media [41], grown at 25°C with continual light. Each inoculation of spore solution was evaluated after 5-7 days to ensure there was germination from >20 spores. After 14-21 days of growth, DNA was extracted from protonema following a modified CTAB/chloroform protocol [40].

### Loci selection and Illumina library preparation

Hypervariable nuclear loci were identified from McDaniel et al. [42]. These loci were amplified via PCR in 15 putative parents (6 males, 9 females) and Sanger sequenced. Discovery and verification of diagnostic SNPs within the putative parents was performed using Geneious v8.1.8, resulting in 5 loci being chosen for use.

Illumina library preparation followed a modification of the Illumina 16S metagenomic protocol (Illumina #15044223 Rev. B) where all loci specific primers have a 33 bp tail added to the 5’ end. This tail contains the binding site of the Illumina sequencing primers and provides a binding site for the indexes (barcodes) and flowcell binding sequences which are added in a 2^nd^ PCR reaction. See Supplemental Materials for more procedural details. The product of the first multiplexed PCR served as the template DNA for the second PCR where each individual’s pool of PCR products was indexed with a 5’ and 3’ index. Custom indexing primers were designed modeled after Illumina Nextera sequence adaptors [43]. The combination of the two indices provided a unique identifier for each individual allowing the pooling and sequencing of several hundred separate libraries in a single Illumina run. The second PCR was carried out that included 1.6 μL product from 1^st^ PCR, run for 10 cycles with a 45 s 55°C annealing temperature. PCR 2 products were visualized and cleaned, then cleaned libraries were quantified and further cleaned. MiSeq 2×250 bp sequencing (Illumina, San Diego, CA, USA) was performed at the University of Florida’s Interdisciplinary Center for Biotechnology Research.

### Genetic data processing and analysis

Raw BCL files, from the MiSeq, had adaptors removed, converted to fastq, and demultiplexed allowing one mismatch in the 5’ and 3’ indexes using Illumina’s bcl2fastq v2.16.0.10 [44]. General patterns observed in FastQC quality plots were used to inform quality trimming parameters. Reads were trimmed using a 10 bp sliding window, with a minimum average quality threshold of 30 using Trim.pl [45]. Trimmed reads were then evaluated again for quality and read length distribution using FastQC. Paired end and singleton reads were assembled against the *Ceratodon purpureus* genome (v0.5) using Bowtie2 v2.2.6 [46]. Since each sample consists of a pool of progeny for each sporophyte, two haplotypes will be present (one corresponding to each parent). Each sample’s BAM file was analyzed using SAMtools to generate two BAM files each containing aligned sequence reads of each corresponding haplotype. SNPs found in the mpileup were called with BCFtools call v1.2 [47], using the multiallelic-caller (-m), ignoring indels (-V), and calling invariant sites. Genomic regions corresponding to the targeted amplicons were extracted from the VCF output using BCFtools filter (-r). The resulting amplicon VCFs were converted to fasta using a custom Perl script that also evaluated read depth at every position, if read depth dropped below 25 for a given position the script would return an N in the fasta sequence file, indicating the absence of sufficient sequencing data to accurately call the nucleotide at that position. Each amplicon’s sequence file was combined with the Sanger sequenced data from putative parents and aligned using MAFFT implemented in Geneious v8.1.8 (BioMatters Ltd, Auckland, New Zealand). The resulting alignments were clustered based on pairwise sequence similarity and every individual’s two haplotype sequences were assigned to a known or unknown parent. Because each locus was not capable on its own of resolving every parent this process was repeated across all loci, ultimately producing a unique multilocus assignment that when compared to the known parents could identify maternal and paternal contributors to the sporophyte.

### Data analysis

We used a two-way ANOVA to determine the effects of microarthropod treatment, sampling date, and the interaction between these effects on CFR in the *C. purpureus* canopies, with CFR log-transformed, and ANOVA to determine the effects of microarthropod treatment on Fv/Fm. GLM tested the effect of microarthropod treatment on sporophyte counts [48].

In genotyping 325 sporophytes, we found that initially-planted genotypes accounted for 95.7% of the paternal genotypes and 85.8% of the maternal genotypes in our sampled sporophytes. Because in some sporophytes we did not sequence the diagnostic SNP which enabled us to distinguish between female genotypes F3 and F6, we assigned these plants to an F6’ maternal parentage. In all cases where we could distinguish between the two, the maternal parent was F3. We used Chi-square tests to determine whether male genotypes differed in their success at producing sporophytes, whether female genotypes differed in their success, and whether there was variation among male genotypes in fathering sporophytes for each female genotype.

We used ANOVA to determine the effects of microarthropod treatment, paternal genotype, and maternal genotype on the distance of each sporophyte from the center of the mesocosm [49]. For each male genotype, we calculated distribution of distances from the center for sporophytes fathered by the genotypes, and whether the distribution was significantly similar to the normal or lognormal distribution using Kolmogorov–Smirnov tests, testing whether sperm dispersal is similar to plant propagule dispersal with a distribution with positive kurtosis (leptokurtic), and may be modeled with the lognormal distribution [50, 51].

We also used ANOVA to test the effects of microarthropod treatment, paternal genotype, and maternal genotype on offspring characters including sporophyte height, percent of spores that germinated, and number of spores produced.

## Results

### Microarthropods and plant parental genotype affect sporophyte production

A total of 839 sporophytes grew across the eight mesocosms with added microarthropods, significantly more than the 228 sporophytes that grew in mesocosms without added microarthropods (Figure 1A; X^2^=345.64; P<0.0001; N=16 mesocosms). To assess potential effects of microarthropods on moss physiology, we measured moss chlorophyll content and photochemical PSII efficiency (Fv/Fm). The addition of microarthropods did not affect chlorophyll content (reported as Chlorophyll Fluorescence Ratio, CFR) in two sampling periods (F=0.5982; P=0.4458; N=32; mean±SE=0.57±0.05 and 0.58±0.05, respectively). There was no significant interaction between microarthropod treatment and sampling period on chlorophyll content (F=0.18 P=0.68), suggesting that the relative chlorophyll content between the treatments did not change across the seasons, nor did microarthropods affect plant physiology.

**Figure 1.**
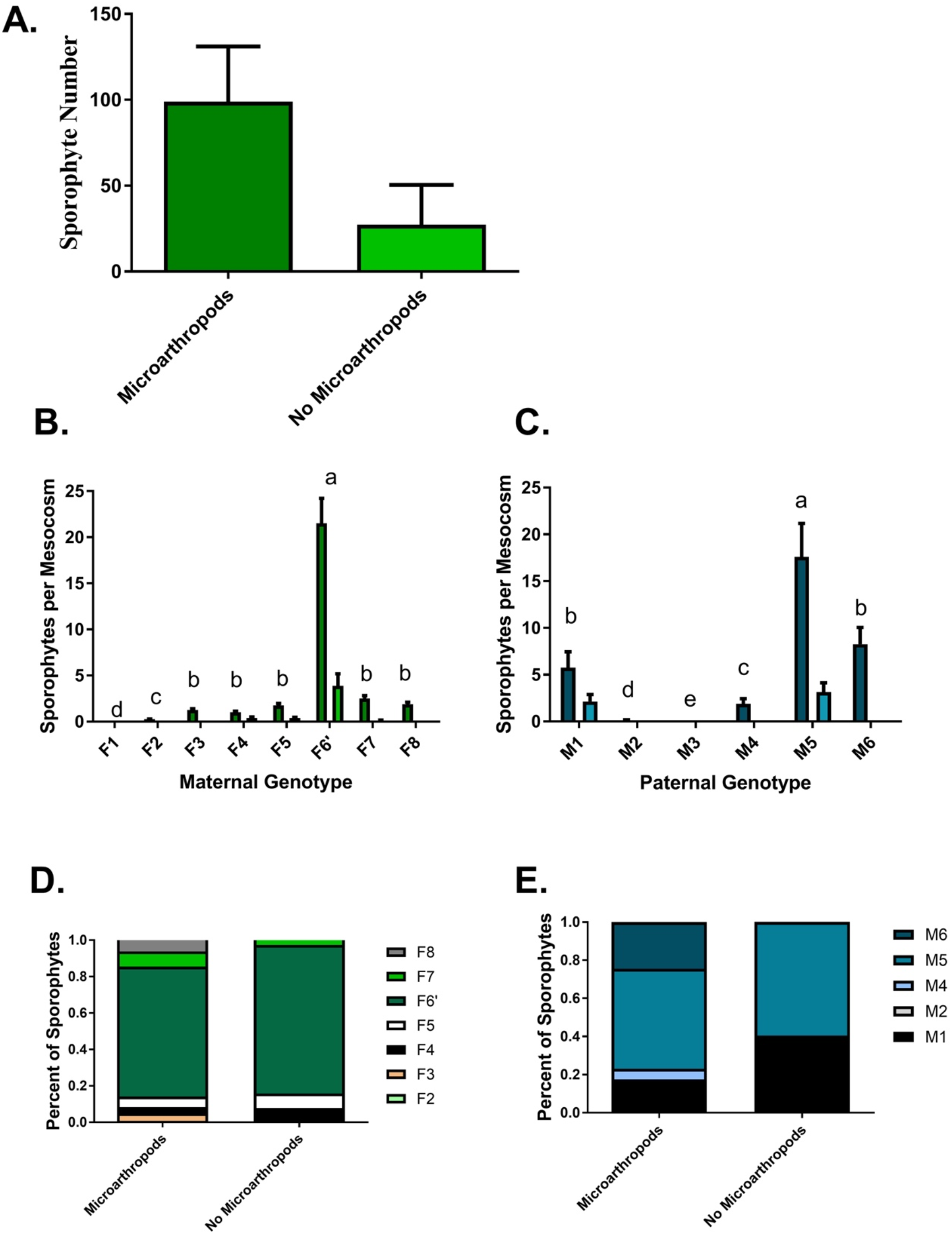
Sporophyte production in the mesocosms. A) Mean (+1 SE) sporophytes produced in mesocosms with and without microarthropods after 16 months (N= 16 mesocosms with 1067 total sporophytes; P<0.0001). Variation in mean (+SE) sporophytes per mesocosm among B) female genotypes (P<0.0001) and C) male genotypes (P<0.0001) produced in treatments with and without microarthropods. Different letters represent significant differences within genotypes for the microarthropod treatments, in which the majority of sporophytes were produced. Comparison of D) maternal genotype and E) paternal genotype numbers in treatments with and without microarthropods (P=0.07 and P<0.0001, respectively).

The moss genotypes used in this experiment had very different fitness, as measured by sporophyte production. Maternal genotype affected sporophyte number (Figures 1B and Table 1; X^2^=276.63; df=4; P<0.0001), with female F6’ producing the most sporophytes. F2 produced only one sporophyte, and F1 produced none. Paternal genotype also had an effect on sporophyte number, with male M5 fathering more sporophytes than the other genotypes in the population, genotypes M2 and M4 fathering fewer, and M3 fathering no sporophytes (Figure 1C; Likelihood Chi-Square=284.41; df=4; P<0.0001). For the majority of females (F3, F5, F6’, F7, and F8), there was significance variation among males in whether they fathered sporophytes with these genotypes (Table 1; df=5; X^2^=14.91, P=0.01; X^2^=14.91, P=0.01; X^2^=21.50, P=0.0007; X^2^=18.73, p=0.002; X^2^=14.91, P=0.01, for the five female genotypes, respectively).

**Table 1.**
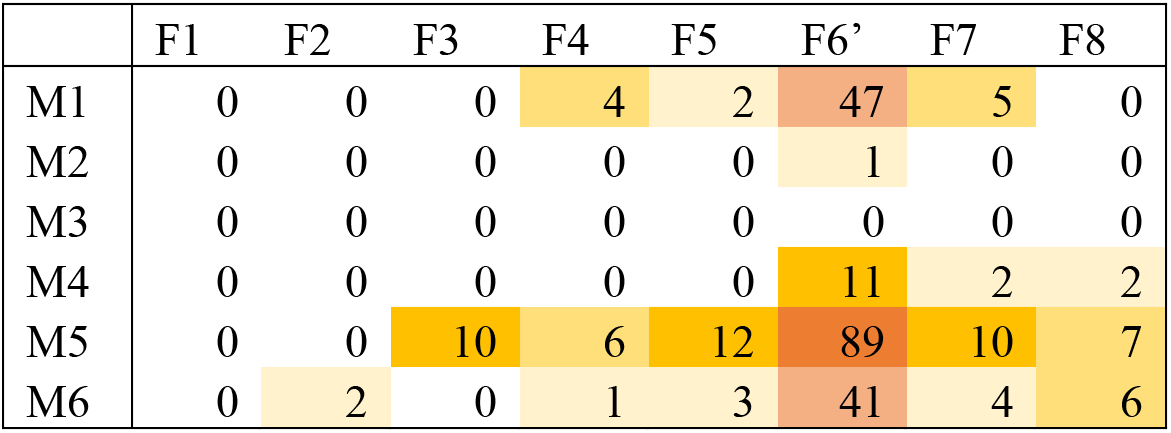
Plant Parental Genotype Affects Fitness. The number of sporophytes produced by each potential pair of male and female genotype. Shading reflects pairs that produced sporophytes, and darker shading reflects pairs that produced higher numbers of sporophytes.

Sporophyte maternal genotype was not significantly affected by microarthropod treatment (X^2^=11.80; P=0.07). However, sporophyte paternal genotype was significantly affected by microarthropod treatment, with mesocosms with microarthropods having sporophytes fathered by five paternal genotypes while mesocosms without microarthropods had sporophytes fathered by only two genotypes (Figure 1E; X^2^=32.08; P<0.0001).

Some male-male competition was evident in the spatial distribution of paternities from the center of a mesocosm (dispersal distance for the sperm from the male genotypes). The distribution of M1 and M4 fertilizations distances was not significantly different from a lognormal distribution (Kologorov’s D=0.059, P=0.15 and Kologorov’s D=0.15, P=0.15, respectively, for goodness of fit to the lognormal distribution). On the other hand, the distributions of the M5 and M6 fertilization distances were significantly different from lognormal (Kologorov’s D=0.10, P=0.01 and Kologorov’s D=0.17, P=0.01, respectively, for goodness of fit to the lognormal distribution). The M1 fertilization distribution had a large positive kurtosis (1.39; indicating a tail away from the center of the pot) while the distribution of M4, M5, and M6 had a negative kurtosis (−0.84, −0.90 and −1.04, respectively, indicating a tail towards the center).

We found that the distance of a sporophyte from the center of the pot (farther from males) was significantly affected by microarthropod treatment (F=20.09; P<0.0001), and paternal genotype (F = 14.50; P<0.0001). Counterintuitively, sporophytes grew farther from the center of the mesocosms (farther from males) in those without microarthropods than with microarthropods (mean±SE distance from the center 13.61±0.30 and 11.19±0.16 cm, respectively), although the distributions were broadly overlapping. Maternal genotype did not affect sporophyte distance from mesocosm center (F=1.33; P=0.2427).

### Haploid parental genotype affects diploid offspring traits

To evaluate the potential to have fitness consequences of different mating choices beyond simply sporophyte production, we also estimated total spore production and average spore germination, two key components of sporophyte fitness, and measured sporophyte height. Sporophyte height was affected by both maternal and paternal haploid genotype (df=6, F=11.25, P<0.0001; and df=4, F=4.59, P=0.0014, respectively), but not microarthropod treatment (df=1, F=0.22, P=0.64). Spore number also was affected by maternal and paternal haploid genotype (df=6, F=4.65, P=0.0004; and df=2, F=4.77, P=0.004, respectively; Figures 3A and 3B), but not by microarthropod treatment (df=1, F=0.35, P=0.55). We found an effect of paternal genotype on spore germination rate (df=3, F=2.90, P=0.05; Figure 3C). Maternal genotype and microarthropod treatment did not influence spore germination rates (df=5, F=1.57, P=0.19; and df=1, F=0.78, P=0.38, respectively; see Supplemental Materials for details).

## Discussion

Here, we show that in large-scale experimental mesocosms, the addition of moss-dwelling microarthropods increases sporophyte formation in the moss *Ceratodon purpureus* by a factor of five (Figure 1A). Importantly, the addition of microarthropods had no apparent effect on chlorophyll fluorescence, suggesting that increased sporophyte formation was unlikely to be due to another factor such as nutrient enhancement resulting in healthier moss.

The addition of microarthropods to our experimental mesocosms also increased the diversity of moss offspring genotypes (Figures 1D and 1E), suggesting that interactions with microarthropods may have a dramatic effect on moss fitness in natural populations. These observations have major implications for quantifying the role of microarthropods in maintaining genetic diversity in natural populations of mosses. Additionally, our results suggest that natural populations of *C. purpureus* which lack commensal arthropods may be sperm-limited, a result previously reported in other moss species [52]. Currently, it is unclear what reward, if any, springtails gain from the behavior, although a moss could provide food sources such as secreted sugars and fatty acids [25, 53, 54], bacteria or fungi [55], or the moss itself [56].

We found significant differences in gamete dispersal distance by male genotype, but it was impractical to map the growth and distribution of each male clone in each mesocosm, therefore we cannot distinguish whether this difference reflects male gametophyte growth or a sperm phenotype. Although spores are the primary moss dispersal stage, selection is likely to favor elevated gamete dispersal, through clonal spreading or sperm production, when mates occur at low density or when levels of inbreeding depression are high. Inbreeding depression is documented in *C. purpureus* [57], meaning that selection for longer distance gamete dispersal might be advantageous.

One remarkable result from this experiment was the abundance of certain male – female genotypic combinations in the resulting sporophytes, but complete lack of other combinations (Table 1). Several factors may explain why all male-female combinations were not equally likely to mate and produce mature sporophytes. Much of the variation is attributable to the fact that some particularly competitive individuals generate more fertilizations (Table 1). However, the absence of combinations of specific competitive male and female genotypes (e.g., F2, F3, or F8 with M1) suggests that some other factor, potentially related to the timing of gamete production [58] or mating compatibility contributes to variation in sporophyte production. Cryptic female choice is possible in mosses because each female makes several archegonia, but only one ever becomes a mature sporophyte. The selective maternal support of only one fertilized egg, or an egg fertilized by a particular sire, provides a plausible explanation for the heterogeneous distribution of offspring genotypes in our mesocosms.

Our data on the spore production from specific male – female combinations also suggest that the genetic variation necessary for parent – offspring conflict is likely to be present in mosses. In mosses all mineral nutrition necessary for sporophyte offspring growth is provided by the maternal gametophyte. Even though sporophytes are photosynthetic for some portion of their lifespan, experimental manipulations show that female gametophytes grow more when dependent sporophytes are removed, suggesting that nurturing the dependent offspring is costly [59]. Haig and Wilczek [60] and Haig [61] proposed a model in which male sex chromosomes evolve to produce more offspring spores by extracting more nutrients from maternal gametophytes. Because mating opportunities may be limited, male fitness scales with the number of spores produced from any single mating. Female fitness, in contrast, is presumably maximized by allocating equally to all of her offspring over the course of her life. Under this model, female sex chromosomes are predicted to evolve defense mechanisms to limit the flow of nutrients across the placenta to the offspring sporophyte. A single female allocating different amounts of energy, as measured by spore production, to offspring sporophytes sired by different males would constitute evidence for genetic conflict over maternal allocation. Females are also variable in their spore production (Figure 2A). Mate choice has shown to influence sporophyte height, which can impact offspring fitness [62, 63], as well as spore number and germination rate. Johnson and Shaw [64] found that male and female haploids had a strong effect on variance of sporophyte fitness in the moss *Sphagnum macrophyllum*. It is possible that females could preferentially support offspring based on some signal of quality, potentially related to sporophyte height or spore production.

**Figure 2.**
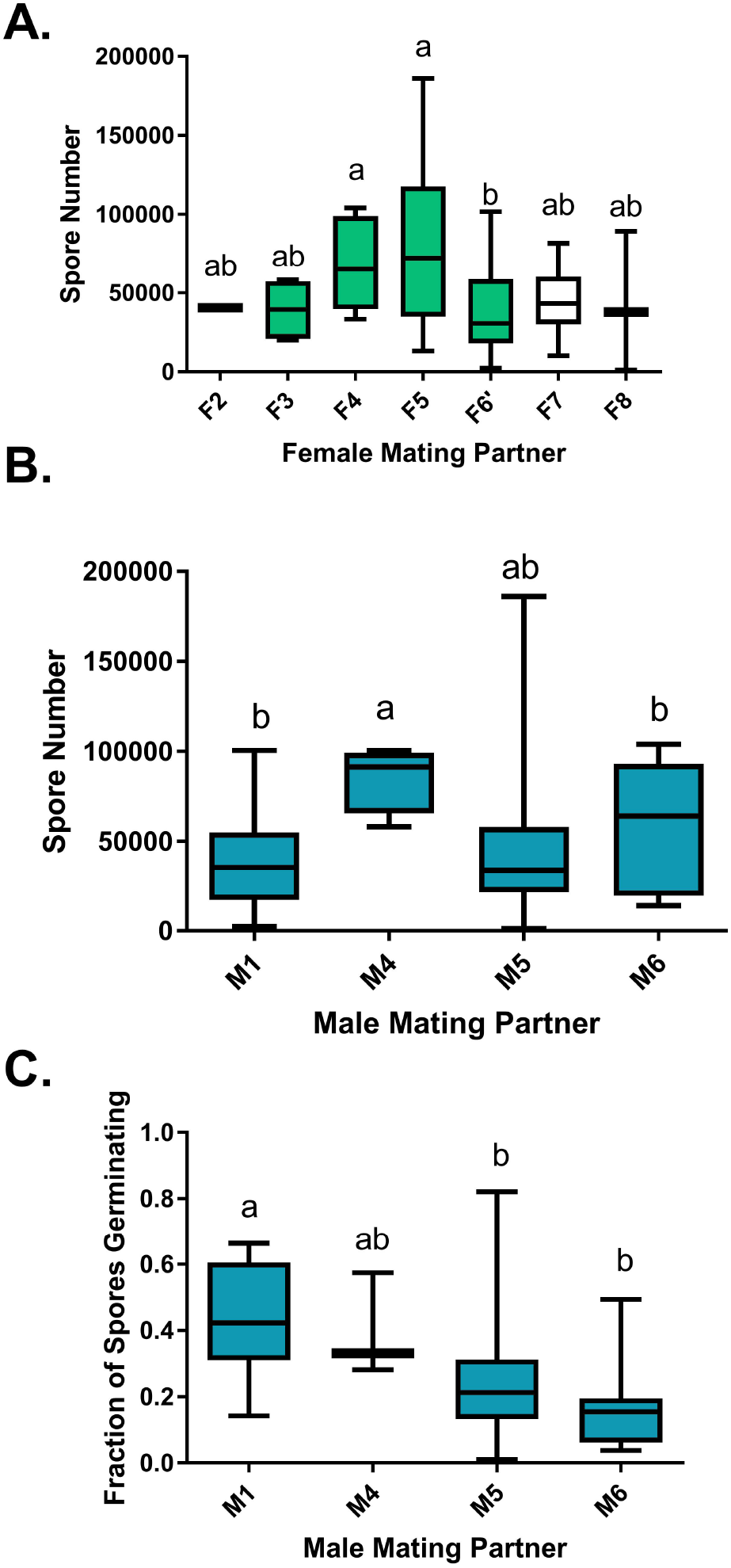
Haploid Parental Genotype Affects Diploid Offspring Traits. Effect of maternal and paternal genotype on offspring traits. A). Spore number produced by each female mating genotype (P=0.0004). B). Spore number produced by each male mating genotype (P=0.004). C) Fraction of spores germinating by each male mating genotype (P=0.05). Maternal genotype did not affect the fraction of germinating spores (P=0.19). Different letters represent significant differences among genotypes.

Collectively these data show that populations of *C. purpureus* are highly polymorphic for components of fitness in both the gametophytic and the sporophytic stages of the life cycle. The presence of sperm-dispersing microarthropods can drive a major increase in fitness, corroborating results suggesting that females are sperm-limited, but may favor certain male genotypes, altering the patterns of male-male competition. In angiosperms, experiments testing female choice and sperm competition suggest that haploid genotypes affect sporophyte fitness [65–69]. Our experiment was underpowered to detect how the presence of microarthropods impacted female mate choice, but our spore production data suggest that such choices could have major consequences for male and female moss fitness. These results clearly highlight the potential dynamic interplay between microarthropods and the maintenance of genetic variation for increased fitness in mosses.

## Conclusions

We found that microarthropods dramatically increase moss sporophyte production, although the sperm dispersal distances were equivalent between mesocosms with microarthropod treatments and the water controls. However, we also found that microarthropods increase the number of moss genotypes that reproduced and differentially influence the fitness of some moss genotypes, resulting in rank-order fitness changes between the treatment and control. These findings suggest that mosses form a commensal relationship with sperm-dispersing microarthropods that may maintain genetic variation for fitness.

## Author Contributions

EES, SME, and TNR designed the experiment. EES conducted the experiment. SFM, ACP, and SBC conducted the genetic analysis. EES and SME analyzed the data and wrote the paper with input from TNR and SFM.

## Acknowledgements

We would like to thank Tera Hinkley, Timea Deakova, Steven Cody Woll, Tina Arredondo, Emily Black, and Trevor Williams for their help in measuring and collecting sporophytes, as well as Linda Taylor for her greenhouse support.

## References

1. Cronberg, N. (2012) Animal-mediated fertilization in bryophytes-parallel or precursor to insect pollination in angiosperms? Lindbergia 35, 76–85.

2. Klavins, S.D. et al. (2005) Coprolites in a Middle Triassic cycad pollen cone: evidence for insect pollination in early cycads?

3. Bao, T. et al. (2019) Pollination of Cretaceous flowers. Proceedings of the National Academy of Sciences 116 (49), 24707–24711.

4. Ashman, T.L. and Majetic, C.J. (2006) Genetic constraints on floral evolution: a review and evaluation of patterns. Heredity 96 (5), 343–352.

5. Patiny, S. (2011) Evolution of Plant-Pollinator Relationships, Cambridge University Press.

6. Waser, N. and Ollerton, J. (2006) Plant-Pollinator Interations, University of Chicago Press.

7. Friedman, J. and Barrett, S.C.H. (2009) Wind of change: new insights on the ecology and evolution of pollination and mating in wind-pollinated plants. Annals of Botany 103 (9), 1515–1527.

8. Harder, L.D. and Wilson, W.G. (1994) Floral evolution and male reproductive success: optimal dispensing schedules for pollen dispersal by animal-pollinated plants. Evolutionary Ecology 8, 542–559.

9. Levitan, D.R. (2005) The distribution of male and female reproductive success in a broadcast spawning marine invertebrate. Integrative and Comparative Biology 45 (5), 848–855.

10. Van Rossum, F. et al. (2011) Fluorescent dye particles as pollen analogues for measuring pollen dispersal in an insect-pollinated forest herb. Oecologia 165 (3), 663–674.

11. Adler, L.S. and Irwin, R.E. (2006) Comparison of pollen transfer dynamics by multiple floral visitors: Experiments with pollen and fluorescent dye. Annals of Botany 97 (1), 141–150.

12. Fenster, C.B. et al. (1996) Fluorescent dye particles are good pollen analogs for hummingbird-pollinated *Silene virginica* (Caryophyllaceae). Canadian Journal of Botany-Revue Canadienne De Botanique 74 (2), 189–193.

13. Sanchez-Robles, J.M. et al. (2014) Effects of tree architecture on pollen dispersal and mating patterns in *Abies pinsapo* Boiss. (Pinaceae). Molecular Ecology 23 (24), 6165–6178.

14. Oddou-Muratorio, S. et al. (2005) Pollen flow in the wildservice tree, Sorbus torminalis (L.) Crantz. II. Pollen dispersal and heterogeneity in mating success inferred from parent-offspring analysis. Molecular Ecology 14 (14), 4441–4452.

15. Hardy, O.J. et al. (2004) Fine-scale genetic structure and gene dispersal in *Centaurea corymbosa* (Asteraceae). II. Correlated paternity within and among sibships. Genetics 168 (3), 1601–1614.

16. Goto, S. et al. (2006) Fat-tailed gene flow in the dioecious canopy tree species *Fraxinus mandshurica* var. *japonica* revealed by microsatellites. Molecular Ecology 15 (10), 2985–2996.

17. Kamm, U. et al. (2009) Frequent long-distance gene flow in a rare temperate forest tree (Sorbus domestica) at the landscape scale. Heredity 103 (6), 476–482.

18. Streiff, R. et al. (1999) Pollen dispersal inferred from paternity analysis in a mixed oak stand of *Quercus robur* L-and Q-petraea (Matt.) Liebl. Molecular Ecology 8 (5), 831–841.

19. Haig, D. (2016) Living together and living apart: the sexual lives of bryophytes. Philosophical Transactions of the Royal Society B-Biological Sciences 371 (1706), 9.

20. Paolillo, D.J., Jr. (1981) The Swimming Sperms of Land Plants. BioScience 31 (5), 367–373.

21. Snäll, T. et al. (2004) Spatial genetic structure in two congeneric epiphytes with different dispersal strategies analysed by three different methods. Molecular Ecology 13 (8), 2109–2119.

22. Jiménez, A. et al. (2010) Microsatellites reveal substantial among-population genetic differentiation and strong inbreeding in the relict fern Dryopteris aemula. Annals of Botany 106 (1), 149–155.

23. Goffinet, B. (2008) Bryophyte biology, Cambridge University Press.

24. Glime, J. (2007) Bryophyte ecology. Ebook sponsored by Michigan Technological University and the International Association of Bryologists. http://www.bryoecol.mtu.edu/. Accessed 20, 2010.

25. Cronberg, N. et al. (2006) Microarthropods mediate sperm transfer in mosses. Science 313 (5791), 1255–1255.

26. Rosenstiel, T.N. et al. (2012) Sex-specific volatile compounds influence microarthropod-mediated fertilization of moss. Nature 481, 431–433.

27. Andrew, N.R. et al. (2003) Variation in invertebrate-bryophyte community structure at different spatial scales along altitudinal gradients. Journal of Biogeography 30, 731–746.

28. Gerson, U. (1969) Moss-arthropod associations. The Bryologist 72, 495–500.

29. Crum, H. and Anderson, L. (1981) Mosses of Eastern North America.

30. Watson, E.V. and Richards, P.R. (1968) British Mosses and Liverworts, Cambridge University Press.

31. Scott, G.A.M. and Stone, I.G. (1976) The Mosses of Southern Australia, Academic Press.

32. Ochyra, R. et al. (2008) The Illustrated Moss Flora of Antarctica, Cambridge University Press.

33. Tsuyuzaki, S. et al. (1999) Vegetation structure in gullies developed by the melting of ice wedges along Kolyma River, northern Siberia. Ecological Research 14 (4), 385–391.

34. Macfadyen, A. (1953) Notes on methods for extractions of small soil arthropods. Journal of Animal Ecology 22 (1), 65–77.

35. Walter, D. (1987) Trophic behavior of mycophagous microarthropods. Ecology 68 (1), 226–229.

36. Yanoviak, S.P. et al. (2004) Arthropod assemblages in vegetative vs. humic portions of epiphyte mats in a neotropical cloud forest. Pedobiologia 48 (1), 51–58.

37. Gitelson, A.A. et al. (1999) The chlorophyll fluorescence ratio F-735/F-700 as an accurate measure of the chlorophyll content in plants. Remote Sensing of Environment 69 (3), 296–302.

38. Bilger, W. et al. (1995) Determination of the quatum efficiency of Photosystem II and of nonphotochemical quenching of chlorophyll fluorescence in the field. Oecologia 102 (4), 425–432.

39. Proctor, M.C.F. (2010) Recovery rates of chlorophyll-fluorescence parameters in desiccation-tolerant plants: fitted logistic curves as a versatile and robust source of comparative data. Plant Growth Regulation 62 (3), 233–240.

40. McDaniel, S.F. et al. (2007) A linkage map reveals a complex basis for segregation distortion in an interpopulation cross in the moss *Ceratodon purpureus*. Genetics 176, 2489–2500.

41. Cove, D.J. and Quatrano, R.S. (2006) Agravitropic mutants of the moss Ceratodon purpureus do not complement mutants having a reversed gravitropic response. Plant Cell and Environment 29 (7), 1379–1387.

42. McDaniel, S.F. et al. (2013) Estimating the nucleotide diversity in *Ceratodon purpureus* (Ditrichaceae) from 218 conserved exon-primed, intron-spanning nucelar loci. Applications in Plant Sciences 1 (4), 13.

43. Mir, K. et al. (2013) Short barcodes for Next Generation Sequencing. Plos One 8 (12), 8.

44. Andrews, S. (2010) FastQC: A quality control tool for high throughput sequence data. http://www.bioinformatics.babraham.ac.uk/projects/FastQC/. (accessed).

45. Nik Joshi (2010) Trim.pl. UC Davis Genomics Center https://github.com/LJI-Bioinformatics/HLATyphon/blob/master/01.Pre_Processing/trim.pl. (accessed).

46. Langmead, B. and Salzberg, S. (2012) Fast gapped-read alignment with Bowtie 2. Nature Methods 9 (4), 357–359.

47. Li, H. et al. (2009) The Sequence Alignment/Map format and SAMtools. Bioinformatics 25 (16), 2078–9.

48. O’Hara, R.B. and Kotze, D.J. (2010) Do not log-transform count data. Methods in Ecology and Evolution 1 (2), 118–122.

49. SAS Institute, JMP for Windows. 13.0.0, SAS Institute, Cary, N.C., 2016.

50. Hintze, C. et al. (2013) D-3: The Dispersal and Diaspore Database - Baseline data and statistics on seed dispersal. Perspectives in Plant Ecology Evolution and Systematics 15 (3), 180–192.

51. Shaw, M.W. et al. (2006) Assembling spatially explicit landscape models of pollen and spore dispersal by wind for risk assessment. Proceedings of the Royal Society B-Biological Sciences 273 (1594), 1705–1713.

52. Bisang, I. et al. (2004) Mate limited reproductive success in two dioicous mosses. Oikos 104 (2), 291–298.

53. Renzaglia, K.S. and Garbary, D.J. (2001) Motile gametes of land plants: Diversity, development, and evolution. Critical Reviews in Plant Sciences 20 (2), 107–213.

54. Paolillo, D. (1979) Lipids of the sperm masses of three mosses. Bryologist 82, 93–96.

55. Balkan, M., Sex-specific fungal communities of the dioicous moss Ceratodon purpureus, Portland State University, USA, 2016.

56. Hågvar, S. (2012) Primary succession in glacier forelands: how small animals conquer new land around melting glaciers. International perspectives on global environmental change 151172.

57. Taylor, P.J. et al. (2007) Sporophytic inbreeding depression in mosses occurs in a species with separate sexes but not in a species with combined sexes. American Journal of Botany 94, 1853–1859.

58. Hendry, A.P. and Day, T. (2005) Population structure attributable to reproductive time: isolation by time and adaptation by time. Molecular Ecology 14 (4), 901–916.

59. Stark, L.R. et al. (2009) An experimental demonstration of the cost of sex and a potential resource limitation on reproduction in the moss *Pterygoneurum* (Pottiaceae). American Journal of Botany 96 (9), 1712–1721.

60. Haig, D. and Wilczek, A. (2006) Sexual conflict and the alternation of haploid and diploid generations. Philosophical Transactions of the Royal Society B: Biological Sciences 361, 335–343.

61. Haig, D. (2013) Kin conflict in seed development: an interdependent but fractious collective. In Annual Review of Cell and Developmental Biology, Vol 29 (Schekman, R. ed), pp. 189–211, Annual Reviews.

62. Budke, J.M. et al. (2013) Dehydration protection provided by a maternal cuticle improves offspring fitness in the moss Funaria hygrometrica. Annals of botany 111 (5), 781–789.

63. Budke, J.M. and Goffinet, B. (2016) Comparative cuticle development reveals taller sporophytes are covered by thicker calyptra cuticles in mosses. Frontiers in plant science 7, 832.

64. Johnson, M.G. and Shaw, A.J. (2016) The effects of quantitative fecundity in the haploid stage on reproductive success and diploid fitness in the aquatic peat moss *Sphagnum macrophyllum*. Heredity 116 (6), 523–530.

65. Skogsmyr, I. and Lankinen, A. (2000) Potential selection for female choice in *Viola tricolor*. Evolutionary Ecology Research 2 (8), 965–979.

66. Waser, N.M. and Price, M.V. (1993) Crossing distance effects on prezygotic preformance in plants: an argument for female choice. Oikos 68 (2), 303–308.

67. Mulcahy, D.L. et al. (1983) Pollen competition in a natural population. In Handbook of Experimental Pollination Biology (Jones, C.E. and Little, R.L. eds), pp. 227–232, Van Rostrand Reinhold.

68. Winsor, J.A. et al. (2000) Pollen competition in a natural population of *Cucurbita foetidissima* (Cucurbitaceae). American Journal of Botany 87 (4), 527–532.

69. Valdivia, E.R. et al. (2007) A group-1 grass pollen allergen influences the outcome of pollen competition in maize. Plos One 2 (1), 7.

