## Supplemental Materials for "Microarthropod contributions to fitness variation in the common moss *Ceratodon purpureus*"

#### Loci selection and Illumina library preparation

Hypervariable nuclear loci were identified from McDaniel et al. (2013). These loci were amplified in 15 putative parents (6 males, 9 females) using OneTaq Hot Start 2x Master Mix (New England Biolabs, Ipswich MA, USA) at PCR conditions appropriate for the primers. Products were Sanger sequenced at the University of Florida's Interdisciplinary Center for Biotechnology Research. Discover and verification of diagnostic SNPs within the putative parents was performed using Geneious v8.1.8, resulting in 5 loci being chosen for use. Primers were redesigned in conserved regions for loci with sequences greater than 500 bp (the maximum length of the available paired end Illumina sequencing) or loci with primes having melting temperatures dissimilar to other loci's primers (desirable for multiplexed PCR, see below).

Illumina library preparation followed a modification of the Illumina 16S metagenomic protocol (Illumina #15044223 Rev. B) where all loci specific primers have a specific 33 bp tail added to the 5' end. This tail contains the binding site of the Illumina sequencing primers and provides a binding site for the indexes (barcodes) and flowcell binding sequences which are added in a 2$^{nd}$ PCR reaction. All tailed loci specific primer pairs were then tested extensively to ensure successful amplification of all five loci in a single multiplexed PCR. Optimal PCR results were achieved with the following conditions: 8 µL OneTaq HotStart 2x MasterMix, 0.1 µL of 10 uM primer each for forward and reverse for all locus, 6 µL PCR H2O, 1 µL DNA, run for 35 cycles with a 45 s 50°C annealing temperature. Low primer concentrations were essential to ensure minimal primer dimers and to keep excess unincorporated primers to a minimum (important for the double PCR design). The product of the first multiplexed PCR served as the template DNA for the second PCR where each individual's pool of PCR products was indexed with a 5' and 3' index. Custom indexing primers were designed modeled after Illumina Nextera sequence adaptors (1000000002694 v00 Oligonucleotide Sequences © 2015 Illumina, Inc), with 7 bp index sequences containing no homopolymer runs and 3 bp mismatch between any two indexes (Mir et al. 2013). The combination of the two indices provided a unique identifier for each individual allowing the pooling and sequencing of several hundred separate libraries in a single Illumina run. The second PCR was carried out with the following conditions: 8 µL OneTaq HotStart 2x MasterMix, 0.4 µL of 10 µM 5' and 3' indexing primer, 5.6 µL PCR H2O, 1.6 µL product from 1$^{st}$ PCR, run for 10 cycles with a 45 sec 55°C annealing temperature. All PCR products were visualized on 1.4% agarose gels to ensure successful amplification prior to the pooling of 5 µL of all PCR 2 products followed by cleaning with a Quiagen PCR clean up kit. Cleaned libraries were quantified with a Qubit Fluorometer (Life Technologies, Grand Island, NY USA), further cleaned with 1.5% AmPure xp (Beckman Coulter, Brea, CA, USA), and visualized to ensure removal of all unincorporated adaptors with Agilent 2100 Bioanalyzer (Agilent Technologies Inc, Santa Clara, CA, USA). MiSeq 2x250 bp sequencing (Illumina, San Diego, CA, USA) was performed at the University of Florida's Interdisciplinary Center for Biotechnology Research.

Oligonucleotide sequences © 2016 Illumina, Inc. All rights reserved. Derivative works created by Illumina customers are authorized for use with Illumina instruments and products only. All other uses are strictly prohibited.

**Genetic data processing and analysis**

All data processing and analysis was carried out on the HiPer Gator Research Computing Cluster at the University of Florida. Raw BCL files, from the MiSeq, had adaptors removed, converted to fastq, and demultiplexed allowing one mismatch in the 5' and 3' indexes using Illumina's bcl2fastq v2.16.0.10. Demultiplexed data was visualized for initial base quality using FastQC v0.11.4 (Andrews

2010). General patterns observed in FastQC quality plots were used to inform quality trimming parameters. All reads were split into separate phased read 1 and read 2 files using a custom Perl script. Reads were trimmed using a 10 bp sliding window, with a minimum average quality threshold of 30 using Trim.pl (Nik Joshi 2010). Trimmed reads where then visualized again for quality and read length distribution using FastQC. Paired end and singleton reads were assembled against the *Ceratodon purpureus* genome (v0.5) using Bowtie2 v2.2.6 (Langmead and Salzberg 2012). We found the best mapping performance was achieved using the sensitive-local assembly parameters. Initial SNP discovery was performed using SAMtools mpileup v1.2 (Li et al. 2009) with the max-depth (-d) set to 25000 and minimum mapping quality for an alignment (-q) set to 30. SNPs found in the mpileup were evaluated and called with BCFtools call v1.2 (Li et al. 2009) using the multiallelic-caller (-m) and ignoring indels (-V). Genomic regions corresponding to the targeted amplicons were extracted from the BCFtools call VCF output using BCFtools filter (-r). The resulting amplicon VCFs were converted to Fasta using a custom Perl script that also evaluated read depth at every position, if read depth dropped below 25 for a given position the script would return an N in the fasta sequence file. Each amplicon's sequence file was combined along with Sanger sequenced amplicons from putative parents and aligned using MAFFT implemented in Geneious v8.1.8 (BioMatters Ltd, Auckland, New Zealand). The resulting nucleotide alignment files were converted to numerical genotype calls and filtered to remove invariant sites as well as sites with a restricted minor allele frequency using custom Perl scripts to allow for their use in the maximum likelihood based parentage analysis program Colony2 v2.0.6.1 (Jones and Wang 2010).

### References


Andrews, S. 2010. FastQC: A quality control tool for high throughput sequence data. http://www.bioinformatics.babraham.ac.uk/projects/FastQC/.

Bateman, A. 1948. Intra-sexual selection in *Drosophila*. Heredity **2**:349-368.

Hintze, C., F. Heydel, C. Hoppe, S. Cunze, A. Konig, and O. Tackenberg. 2013. D-3: The Dispersal and Diaspore Database - Baseline data and statistics on seed dispersal. Perspectives in Plant Ecology Evolution and Systematics **15**:180-192.

Johnson, M. G., and A. J. Shaw. 2016. The effects of quantitative fecundity in the haploid stage on reproductive success and diploid fitness in the aquatic peat moss *Sphagnum macrophyllum*. Heredity **116**:523-530.

Jones, O. R., and J. L. Wang. 2010. COLONY: a program for parentage and sibship inference from multilocus genotype data. Molecular Ecology Resources **10**:551-555.

Langmead, B., and S. Salzberg. 2012. Fast gapped-read alignment with Bowtie 2. Nature Methods **9**:357-359.

Li, H., B. Handsaker, A. Wysoker, T. Fennell, J. Ruan, N. Homer, G. Marth, G. Abecasis, R. Durbin, and P. Genome Project Data. 2009. The Sequence Alignment/Map format and SAMtools. Bioinformatics **25**:2078-2079.

McDaniel, S. F., M. J. van Baren, K. S. Jones, A. C. Payton, and R. S. Quatrano. 2013. Estimating the nucleotide diversity in *Ceratodon purpureus* (Ditrichaceae) from 218 conserved exon-primed, intron-spanning nucelar loci. Applications in Plant Sciences **1**:13.

McDaniel, S. F., J. H. Willis, and A. J. Shaw. 2007. A linkage map reveals a complex basis for segregation distortion in an interpopulation cross in the moss *Ceratodon purpureus*. Genetics **176**:2489-2500.

Mir, K., K. Neuhaus, M. Bossert, and S. Schober. 2013. Short barcodes for Next Generation Sequencing. Plos One **8**:8.

Nik Joshi. 2010. Trim.pl. UC Davis Genomics Center
https://github.com/LJI-Bioinformatics/HLATyphon/blob/master/01.Pre_Processing/trim.pl.

O'Hara, R. B., and D. J. Kotze. 2010. Do not log-transform count data. Methods in Ecology and Evolution **1**:118-122.

SAS Institute. 2015. JMP for Windows. Release 12.0.1. Cary, N.C.

Shaw, M. W., T. D. Harwood, M. J. Wilkinson, and L. Elliott. 2006. Assembling spatially explicit landscape models of pollen and spore dispersal by wind for risk assessment. Proceedings of the Royal Society B-Biological Sciences **273**:1705-1713.
